# Pharmacologic Targeting of PDIA1 Inhibits NLRP3 Inflammasome Assembly and Activation

**DOI:** 10.1101/2023.08.21.554183

**Authors:** Jessica D. Rosarda, Caroline R. Stanton, Emily B. Chen, Michael J. Bollong, R. Luke Wiseman

**Author notes:** To whom correspondences should be addressed: R. Luke Wiseman, Department of Molecular Medicine Scripps Research, La Jolla, CA 92037.

## Abstract

The NLRP3 inflammasome is a cytosolic protein complex that regulates innate immune signaling in response to diverse pathogenic insults through the proteolytic processing and secretion of pro-inflammatory cytokines such as IL-1β. Hyperactivation of NLRP3 inflammasome signaling is implicated in the onset and pathogenesis of numerous diseases, motivating the discovery of new strategies to suppress NLRP3 inflammasome activity. We sought to define the potential for the proteostasis regulator AA147 to inhibit the assembly and activation of the NLRP3 inflammasome. AA147 is a pro-drug that is metabolically converted to a reactive metabolite at the endoplasmic reticulum (ER) membrane to covalently modify ER-localized proteins such as protein disulfide isomerases (PDIs). We show that AA147 inhibits NLRP3 inflammasome activity in monocytes and monocyte-derived macrophages through a mechanism involving impaired assembly of the active inflammasome complex. This inhibition is mediated through AA147-dependent covalent modification of PDIA1. Genetic depletion or treatment with other highly selective PDIA1 inhibitors similarly blocks NLRP3 inflammasome assembly and activation. Our results identify PDIA1 as a potential therapeutic target to mitigate NLRP3 inflammasome-mediated pro-inflammatory signaling implicated in etiologically diverse diseases.

## INTRODUCTION

Inflammasomes are multi-protein complexes involved in orchestrating local and systemic innate immune responses to diverse insults.^1–3^ While multiple inflammasome complexes have been identified, activation of the NLR family pyrin domain containing 3 (NLRP3) inflammasome is perhaps the most clearly understood.^3,4^ The NLRP3 inflammasome can be activated by a multitude of pathogen- or damage-associated signals which generally involve both a priming and an activating stimulus.^2,3,5^ The priming signal induces the NF-κB dependent expression of NLRP3 inflammasome signaling components, while a secondary activation signal then initiates oligomerization of individual components into the functional NLRP3 inflammasome complex (**Fig. S1A**).^3,6,7^ Upon assembly and activation, the NLRP3 inflammasome stimulates the maturation and release of pro-inflammatory cytokines such as the highly potent cytokine IL-1β.^1,2,8^ Secreted cytokines then promote inflammatory responses that can neutralize threats, although this can also contribute to inflammation-mediated tissue damage.^9,10^ Pathologic IL-1β signaling caused by NLRP3 inflammasome hyperactivation has been linked to diverse diseases including cryopyrin-associated periodic syndromes (CAPS)^11,12^, type II diabetes mellitus^9,13,14^, as well as cardiac and cerebral ischemia and reperfusion (I/R) injury.^8–10,15,16^

Considering the central importance of the NLRP3 inflammasome in the pathogenesis of disease, therapeutic strategies to inhibit the activation and signaling of this complex has emerged as a highly attractive approach to mitigate inflammation-related pathologies.^7,17–19^ In fact, inhibiting the NLRP3 inflammasome and downstream signaling has been shown to reduce tissue damage and improve outcomes in a variety of animal disease models, as well as in human patients.^10,13,16,19,20^ Monoclonal antibodies that suppress IL-1β signaling downstream of inflammasome activation are clinically approved to treat several inflammatory diseases and have shown benefits in numerous animal models of inflammatory disorders.^10,13,20,21^ Further, numerous pharmacologic strategies have been developed to inhibit NLRP3 inflammasome assembly and activation.^18^ Compounds like MCC950 have been shown to bind the ATP binding pocket of NLRP3 to inhibit inflammasome assembly and mitigate pathologic inflammasome signaling in numerous mouse models of inflammatory disease.^15,22–25^ Other compounds that bind the NLRP3 ATP binding site, such as OLT1177, have also been tested in clinical trials for inflammatory diseases such as gout and osteoarthritis.^26,27^ Despite the promise of these approaches, no direct NLRP3 inflammasome inhibitor is currently approved as a treatment, necessitating the continued development of new pharmacologic strategies to block NLRP3 inflammasome signaling.

Intriguingly, numerous electrophilic compounds have also been found to covalently modify cysteine residues within the NLRP3 inflammasome and inhibit complex assembly. The anti-inflammatory compound oridonin inhibits NLPR3 inflammasome activity through a mechanism predicted to involve selective modification of NLRP3^Cys279^.^28^ Similarly, compound RRx-001, which was originally identified as an anticancer compound, is predicted to inhibit inflammasome assembly through covalent modification of NLRP3^Cys409^.^29^ Further, itaconate, an immunomodulatory metabolite, is predicted to block inflammasome assembly and activity in mice through modification of NLRP3^Cys548^.^30^ More recent screens have identified additional electrophilic compounds that target multiple cysteines residues on NLRP3 to inhibit inflammasome assembly and activity, indicating that NLRP3 is a major electrophilic sensor within the cell.^31^

We previously identified compound AA147 as a highly-selective regulator of endoplasmic reticulum (ER) proteostasis.^32^ We showed that the 2-amino-p-cresol moiety of AA147 is metabolically activated at the ER membrane to a reactive electrophile that selectively modifies proteins localized near the ER membrane, such as protein disulfide isomerases (PDIs).^33^ AA147-dependent PDI modification can both directly influence ER proteostasis for destabilized proteins such as immunoglobulin light chains^34^ and induce adaptive ER proteostasis remodeling through activation of the ATF6 signaling arm of the unfolded protein response.^32^ Previous results suggest that the NLRP3 inflammasome assembles at the ER membrane.^35^ Considering the electrophilic sensing activity of NLRP3^31^, we predicted that local, metabolic activation of AA147 to a reactive electrophile could potentially inhibit NLRP3 inflammasome assembly and activity through direct targeting of NLRP3 cysteine residues.^31,36^ Supporting this prediction, we previously showed that AA147 covalently modifies the electrophilic sensor KEAP1 to activate NRF2 signaling in neuronal cell models through a similar mechanism.^36^

Here, we found that AA147 is a potent and effective inhibitor of NLRP3 inflammasome assembly and activation. However, this inhibition is independent of both AA147-dependent ATF6 activation. Instead, AA147 blocks inflammasome assembly and activation through a mechanism involving covalent modification of PDIA1 – a highly abundant ER PDI. Genetic depletion of *PDIA1* mimics AA147-depenent inflammasome inhibition. We also demonstrate that highly selective PDIA1 inhibitors show potent activity against NLRP3 inflammasome activation. These results identify PDIA1 as a novel target for inhibiting inflammasome signaling that could be potentially exploited for the continued development of next generation inflammasome inhibitors.

## RESULTS

### AA147 reduces NLRP3 assembly and activity

We initially sought to define the potential for the metabolically activated pro-drug AA147 to inhibit NLRP3 inflammasome assembly. We employed a recently described, high content imaging-based screening assay of inflammasome assembly that utilizes the human monocyte cell line THP1 expressing the inflammasome subunit ASC fluorescently tagged with GFP (ASC-GFP) under control of an NFκB-regulated promoter.^31^ Inflammasome assembly can be monitored in these cells by following the formation of ASC-GFP puncta, or ‘specks’, that accumulate in response to treatment with both a priming stimulus (e.g., LPS) and an activating stimulus (e.g., nigericin, ATP). Treatment with both LPS and nigericin robustly increases ASC-GFP speck formation in undifferentiated THP1 cells (**Fig. 1A,B**). Pre-treatment with the known NLRP3 inflammasome inhibitor MCC950^23^ inhibited ASC-GFP speck formation. Interestingly, pre-treatment with AA147 similarly blocked ASC-GFP speck formation in undifferentiated THP1 cells, indicating that this compound inhibits inflammasome assembly.

**Figure 1.**
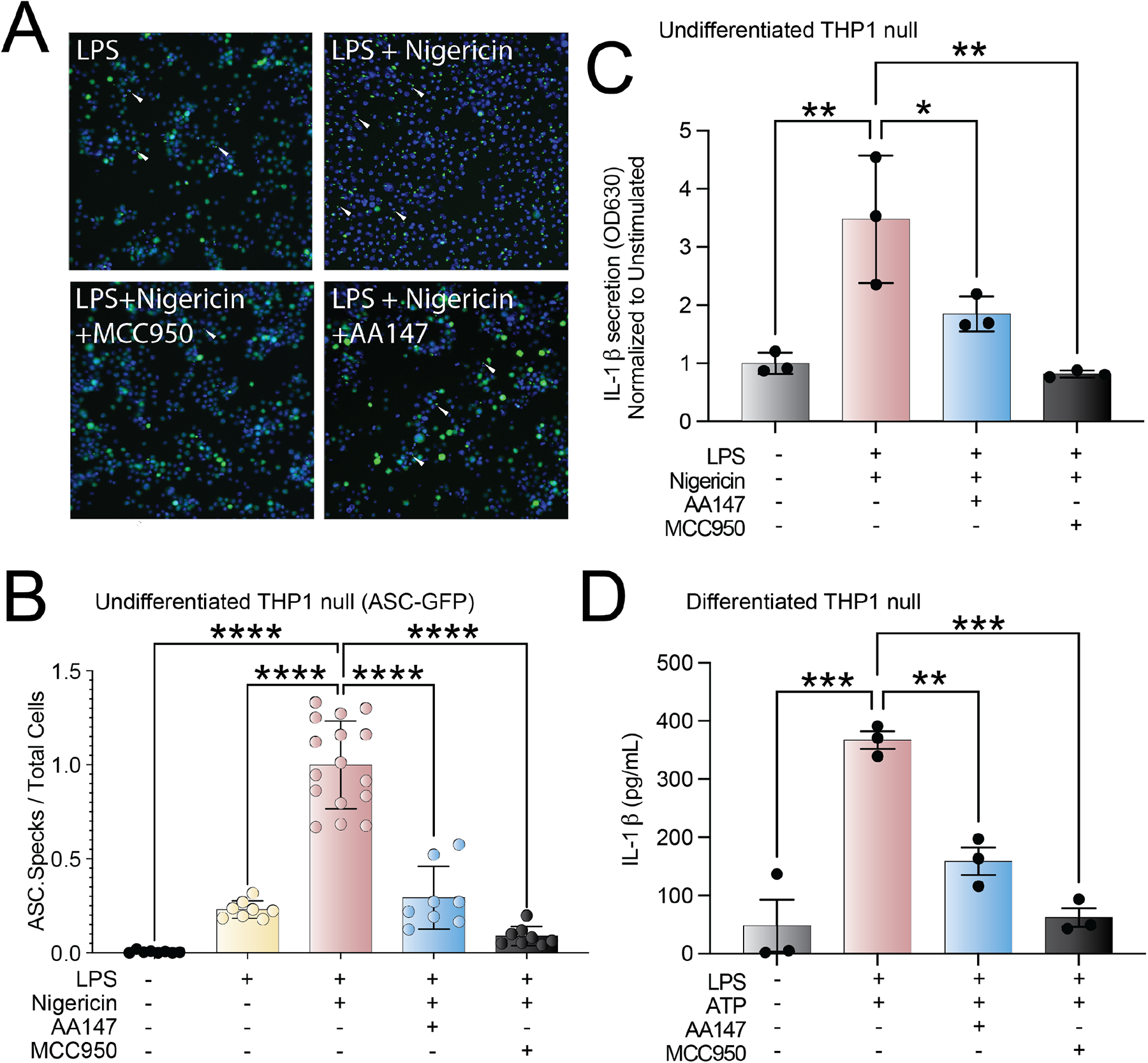
AA147 reduces NLRP3 inflammasome assembly and IL-1β secretion. **A**. Representative fluorescent microscopy images of ASC-GFP cells untreated, treated with LPS (1µg/mL) for 16 h ± nigericin (1 µM) 3 hours and in the presence or absence of AA147 (10 µM). ASC-GFP specks are indicated with white arrows. **B**. Quantification of images shown in (A). **** p <0.0001 using a Brown-Forsythe and Welch ANOVA test with Dunnett-T3 correction for multiple comparisons. Error bars show SD. **C.** Quantification of secreted bioactive IL-1β assessed using the SEAP Quanti-blue assay. HEK Blue IL-1β cells were treated with media conditioned on THP1 monocytes pre-treated with AA147 (10 µM) or vehicle for 16 h and stimulated with LPS (1µg/mL) and nigericin (1 µM) for 24 h relative to an unstimulated, vehicle treated control. **** p <0.0001, *** p < 0.001, ** p<0.01 for one-way ANOVA test with Dunnett correction for multiple comparisons. Error bars show SEM for n=3. **D.** ELISA quantifications of total IL-1β detected in media conditioned on THP1 monocyte-derived macrophages pre-treated with AA147 (10 µM) or vehicle for 16 h then stimulated with LPS (1µg/mL) for 3 h and incubated with ATP (5 mM) for 24 h. *** p < 0.001, ** p<0.01 for one-way ANOVA test with Dunnett correction for multiple comparisons. Error bars show SEM for n=3.

Inflammasome activation leads to the increased secretion of the pro-inflammatory cytokine IL-1β.^7^ Thus, we sought to determine the extent to which AA147 blocks this activity downstream of inflammasome assembly. Initially, we monitored secretion of IL-1β from monocyte THP1 cells using HEK-Blue IL-1β reporter cells (Invivogen) – a cell line that expresses the IL-1β receptor and an NFκB-responsive secreted alkaline phosphatase (SEAP) reporter. Treatment of HEK-Blue IL-1β cells with conditioned media prepared on THP1 cells stimulated with LPS and nigericin increased secretion of SEAP, reflecting increased IL-1β secretion (**Fig. 1C**). Treatment of THP-1 cells with MCC950 blocked this effect, confirming that the increase in SEAP corresponds to NLRP3 inflammasome-dependent increases in IL-1β secretion from THP1 cells. AA147 similarly blocked IL-1β secretion from undifferentiated THP1 cells stimulated with LPS and nigericin (**Fig. 1C**). Similar results were observed in THP1 monocyte-derived macrophages stimulated with LPS and ATP, as measured by both the HEK-Blue IL-1β reporter assay (**Fig. S1B**) and an enzyme-linked immunoassay (ELISA) of conditioned media (**Fig. 1D**). Further, AA147 reduced IL-1β in conditioned media prepared on RAW264.7 mouse macrophages stimulated with LPS and ATP, as measured by immunoblotting (**Fig. S1C**). These results indicate that AA147 reduces NLRP3 inflammasome assembly and activity.

### AA147-dependent inhibition of inflammasome activation is independent of ATF6 activation or NFκB inhibition

AA147 was originally identified as an activator of the ATF6 signaling arm of the unfolded protein response.^32^ Thus, we sought to determine the potential for AA147 to inhibit inflammasome activity through an ATF6-dependent mechanism. Surprisingly, we did not observe significant increases of the ATF6-target BIP in immunoblots prepared from THP1 cells treated for 16 h with AA147, indicating that this compound did not significantly increase ATF6 signaling in these cells (**Fig. S2A**). Further, co-treatment with the ATF6 inhibitors Ceapin-A7^37^ or the site 1 protease inhibitor PF-429242^38^ did not decrease AA147-dependent reductions of IL-1β secretion in monocyte THP1 cells stimulated with LPS and ATP (**Fig. 2A,B**). These results indicate that AA147 does not reduce NLRP3 inflammasome-dependent IL-1β secretion through an ATF6-dependent mechanism.

**Figure 2.**
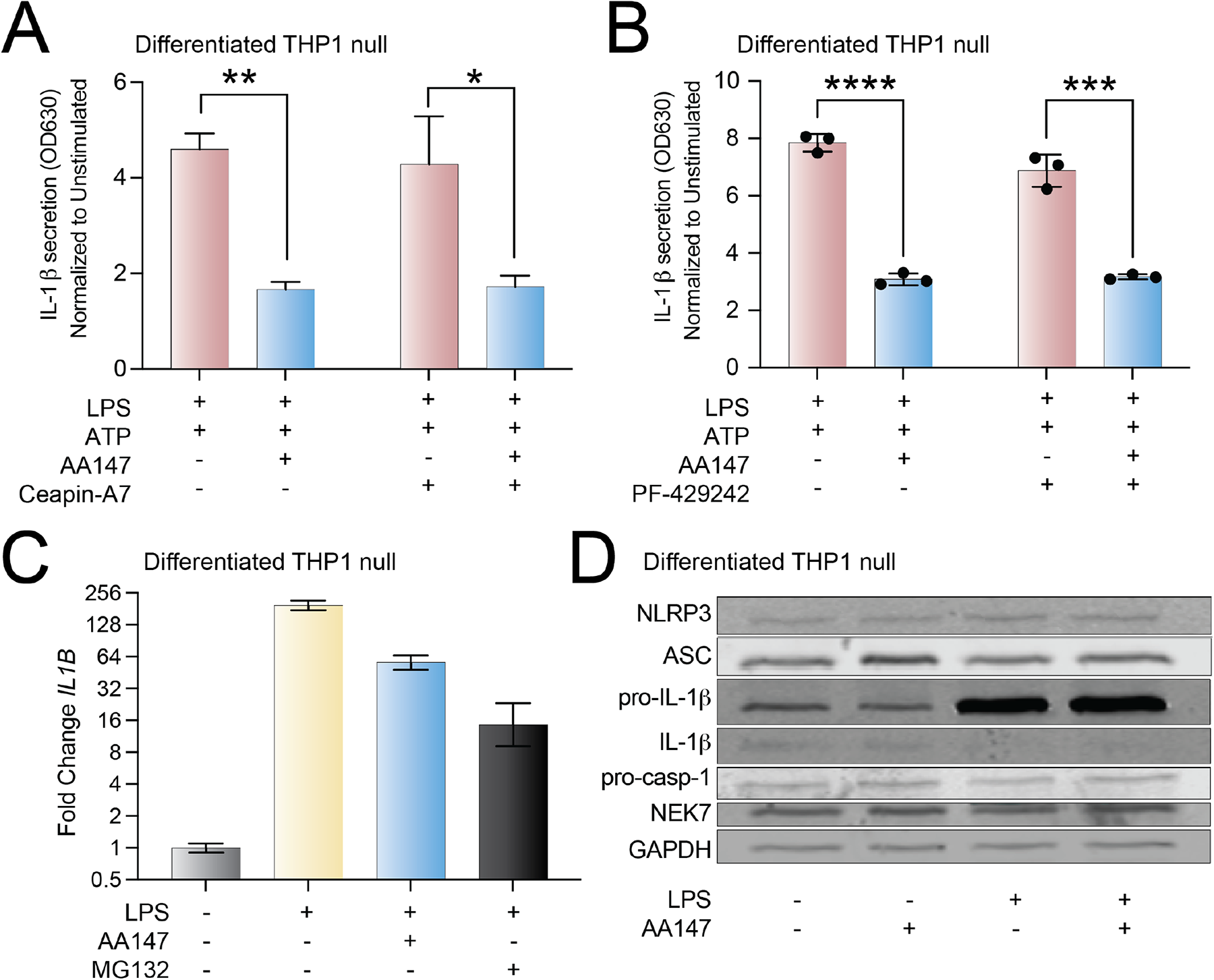
AA147-dependent inhibition of NLRP3 inflammasomes is independent of ATF6 activation. **A**. Quantification of secreted bioactive IL-1β assessed using the SEAP Quanti-blue assay. HEK Blue IL-1β cells were treated with media conditioned on THP1 monocyte-derived macrophages pre-treated with AA147 (10 µM) or vehicle for 16 h in the presence or absence of the ATF6 inhibitor Ceapin-A7 (CP7; 6 µM) and subsequently stimulated for 3 h with LPS (1µg/mL) and 24 h with ATP (5 mM). Results are relative to unstimulated, vehicle treated controls. *** p < 0.001, **** p <0.0001 with multiple unpaired two-tailed t-tests. Error bars show SEM for n=3. **B**. Quantification of secreted bioactive IL-1β assessed using the SEAP Quanti-blue assay. HEK Blue IL-1β cells were treated with media conditioned on THP1 monocyte-derived macrophages pre-treated with AA147 (10 µM) or vehicle for 16 h in the presence or absence of the ATF6 inhibitor PF429242 (10 µM) and subsequently stimulated for 3 h with LPS (1µg/mL) and 24 h with ATP (5 mM). Results are relative to unstimulated, vehicle treated controls. *** p < 0.001, **** p <0.0001 with multiple unpaired two-tailed t-tests. Error bars show SEM for n=3. **C**. mRNA expression measured using qPCR, of the pro-inflammatory cytokine *IL1B* in THP1 monocyte-derived macrophages treated for 16 h with vehicle or AA147 (10 µM) and subsequently stimulated with LPS (1µg/mL) for 3 h. Error bars show 95% CI. **D**. Immunoblot of inflammasome components in THP1 monocyte-derived macrophages pre-treated with vehicle or AA147 (10 µM) for 16 h and subsequently stimulated with vehicle or LPS (1µg/mL) for 3 h.

AA147 could also influence inflammasome activation through suppression of NFκB-dependent expression of inflammasome components and pro-IL-1β. AA147 modestly reduced expression of *IL1B* in THP1 monocyte derived macrophages stimulated with LPS (**Fig. 2C**), although this reduction was not detectable in naïve THP1 cells which showed a less dynamic response to LPS (**Fig. S2B**). Further, AA147 did not impact the expression of other NLRP3 inflammasome subunits in differentiated THP1 cells (e.g., *ASC*, *NLRP3*, *CASP1*)(**Fig. S2C**). We also did not observe changes in the protein levels for NLRP3 subunits or pro-IL-1β in differentiated THP1 cells stimulated with LPS and treated with AA147 (**Fig. 2D**). This suggests that AA147 does not significantly impact expression of inflammasome subunits or pro-IL-1β in these cells.

### Genetic depletion of PDIA1 inhibits inflammasome assembly and activation

AA147 activates ATF6 through a mechanism involving metabolic activation and direct covalent modification of a subset of ER protein disulfide isomerases (PDIs).^33^ Previous results showed that AA147-dependent inhibition of PDIs could promote adaptive ER remodeling independent of ATF6 to influence secretion of destabilized, amyloidogenic immunoglobulin light chains.^34^ Thus, we sought to determine if NLRP3 inflammasome activation and assembly could also be explained by AA147-dependent PDI inhibition. We used an alkyne-containing analog of AA147 (AA147^alk^)^33^ to confirm that AA147 covalently modified PDIs in monocyte THP1 cells (**Fig. S3A**). We then shRNA-depleted the predominant PDIs modified by AA147, including *PDIA1*, *PDIA3*, *PDIA4*, *PDIA5*, and *PDIA6* ^33,34^, in THP1 cells stimulated with LPS and ATP or nigericin and then monitored IL-1β secretion using the HEK-Blue IL-1β reporter assay. We confirmed efficient knockdown of individual PDIs by qPCR (**Fig. S3B**). While depletion of all tested PDIs show modest inhibition of IL-1β secretion from THP1 cells stimulated with LPS and either nigericin or ATP, the most significant effects were observed upon knockdown of *PDIA1* and *PDIA5* (**Fig. 3A**). We specifically focused on PDIA1, as this PDI is highly modified by AA147 across multiple cell types, whereas PDIA5 has not been identified as a robust target across cell types.^33,34,36^ Additionally, CRISPR-mediated ablation of PDIA1 diminishes secretion of pro-inflammatory cytokines in other cell types, implicating PDIA1 as a potential regulator of inflammatory signaling.^39^

**Figure 3.**
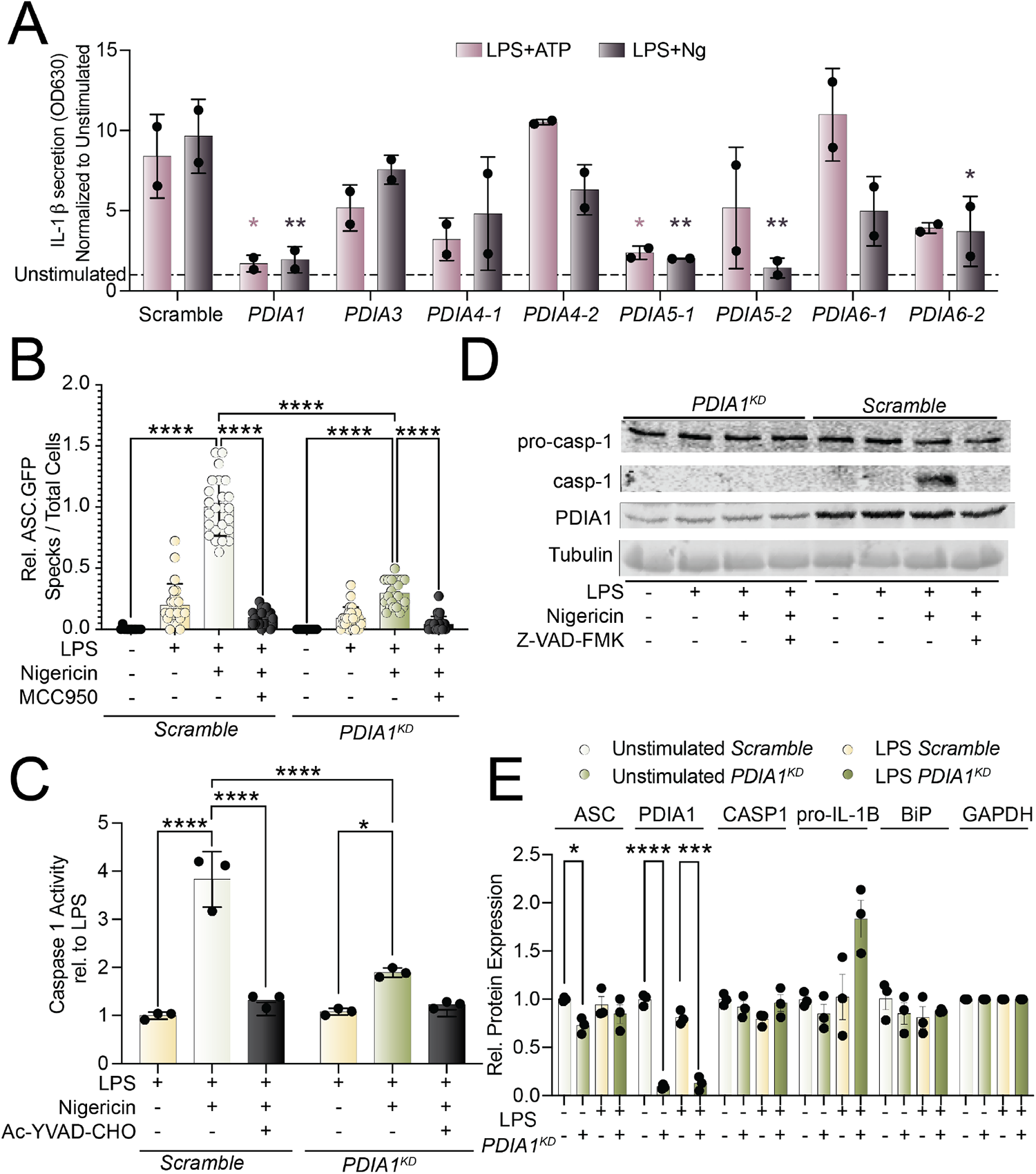
*PDIA1* depletion inhibits NLRP3 inflammasome activation. **A.** Quantification of secreted bioactive IL-1β assessed using the SEAP Quanti-blue assay. HEK Blue IL-1β cells were treated with media conditioned on THP1 monocytes stably expressing shRNAs of the respective target PDI. were stimulated for 3 h with LPS (1µg/mL) and 24 h with ATP (5 mM) or nigericin (1 μM). * p < 0.05, *** p < 0.001, **** p <0.0001 two-way ANOVA test with Dunnett correction for multiple comparisons. Error bars show SD. **B**. Relative number of ASC-GFP specks per cell detected using fluorescent microscopy in undifferentiated THP1 ASC-GFP monocyte cells stably expressing either a scramble shRNA or a PDIA1 shRNA (PDIA1^KD^). Cells were stimulated with LPS (1µg/mL) for 16 h and nigericin (5 µM) for 3 h. Quantification is relative to the overall average number of specks / total cells across all wells in LPS (1µg/mL) and nigericin (5 µM) stimulated wells. **** p <0.0001 using a Brown-Forsythe and Welch ANOVA test with Dunnett-T3 correction for multiple comparisons. Error bars show SD. **C.** Caspase-1 activity assessed using Caspase-1 Glo assay from PDIA1^KD^ or scramble THP1 monocytes stimulated for 16 h with LPS (1µg/mL) and 45 min nigericin (1 μM). Error bars show SD; each dot represents an individual replicate relative to LPS. **** p <0.0001 using a two-way ANOVA. **D.** Immunoblot of cell lysates from PDIA1^KD^ or scramble THP1 cells stimulated with LPS (1µg/mL) for 3 h and nigericin (5 µM) for 1 h. ZVAD-VAD-FMK (50 µM) was used as a positive control. **E**. Quantification of protein levels measured using immunoblots from cell lysates from *PDIA1^KD^* or scramble THP1 cells stimulated with LPS (1µg/mL) for 3 h. * p< 0.05, *** p <0.001. **** p<0.0001 using multiple unpaired two-tailed t-tests. Error bars show SEM for n = 3.

We confirmed that PDIA1 protein was reduced in THP1 cells stably depleted of *PDIA1* (**Fig. S3C**). Importantly, we also showed that *PDIA1*-depletion did not increase expression of the UPR target gene BiP, indicating that reduced activity of PDIA1 does not globally induce ER stress in these cells. As with AA147 pretreatment, *PDIA1* depletion reduced ASC-GFP speck formation in THP1 cells stimulated with LPS and nigericin (**Fig. 3B**). Further, we found that *PDIA1* depletion inhibited autoproteolysis of caspase 1 (**Fig. 3C,D**) – another marker of inflammasome assembly and activation.^7,40^ *PDIA1* depletion showed a modest reduction in *IL1B* expression in LPS stimulated THP1 monocytes, although no effect was observed for the NLRP3 inflammasome subunits *NLRP3* or *ASC* (**Fig. S3C**). However, we did not observe significant reductions in pro-IL-1β in THP1 cells depleted of *PDIA1* (**Fig. 3E**). These results indicate that *PDIA1* depletion inhibits NLRP3 inflammasome assembly and activation through a mechanism analogous to that observed with AA147.

### Selective, pharmacologic PDIA1 inhibitors block inflammasome assembly and activation

The above results suggest that AA147 inhibits NLRP3 assembly and activation through covalent targeting of PDIs, most notably PDIA1. This suggests that other PDIA1 inhibitors should similarly inhibit NLRP3 inflammasome assembly and activation. To test this, we monitored ASC-GFP speck formation in monocyte THP1 cells treated with the highly selective, covalent PDIA1 inhibitor KSC-34^41^ and stimulated with LPS and nigericin. KSC-34 contains an alkyne moiety, allowing us to confirm efficient engagement of PDIA1 by this compound (**Fig. 4A, B**). Treatment with KSC-34 significantly reduced ASC-GFP speck formation induced by LPS and nigericin (**Fig. 4C**). Further, KSC-34 reduced IL-1β secretion in THP1 cells stimulated with LPS and nigericin, as measured by HEK-Blue IL-1β reporter assay (**Fig. S4A**). Similar results were observed in THP1 cells stimulated with LPS and ATP (**Fig. 4A**). The structurally analogous PDIA1 inhibitor RB-11-CA^41^ also demonstrated reduced IL-1β secretion from monocyte THP1 cells (**Fig. 4A,B,E**). Further, we confirmed that these two PDIA1 inhibitors did not directly label NLRP3, unlike the recently identified NLRP3 covalent modifier P207-9174 (**Fig. S4B**).^31^ Lastly, we demonstrated that neither AA147 nor KSC-34 showed additional reduction in IL-1β secretion, measured by HEK-IL-1β reporter assay, from THP1 cells depleted of *PDIA1* and stimulated with LPS and nigericin (**Fig. S4C**). This indicates that these inhibit inflammasome activity through a *PDIA1*-dependent mechanism.

**Figure 4.**
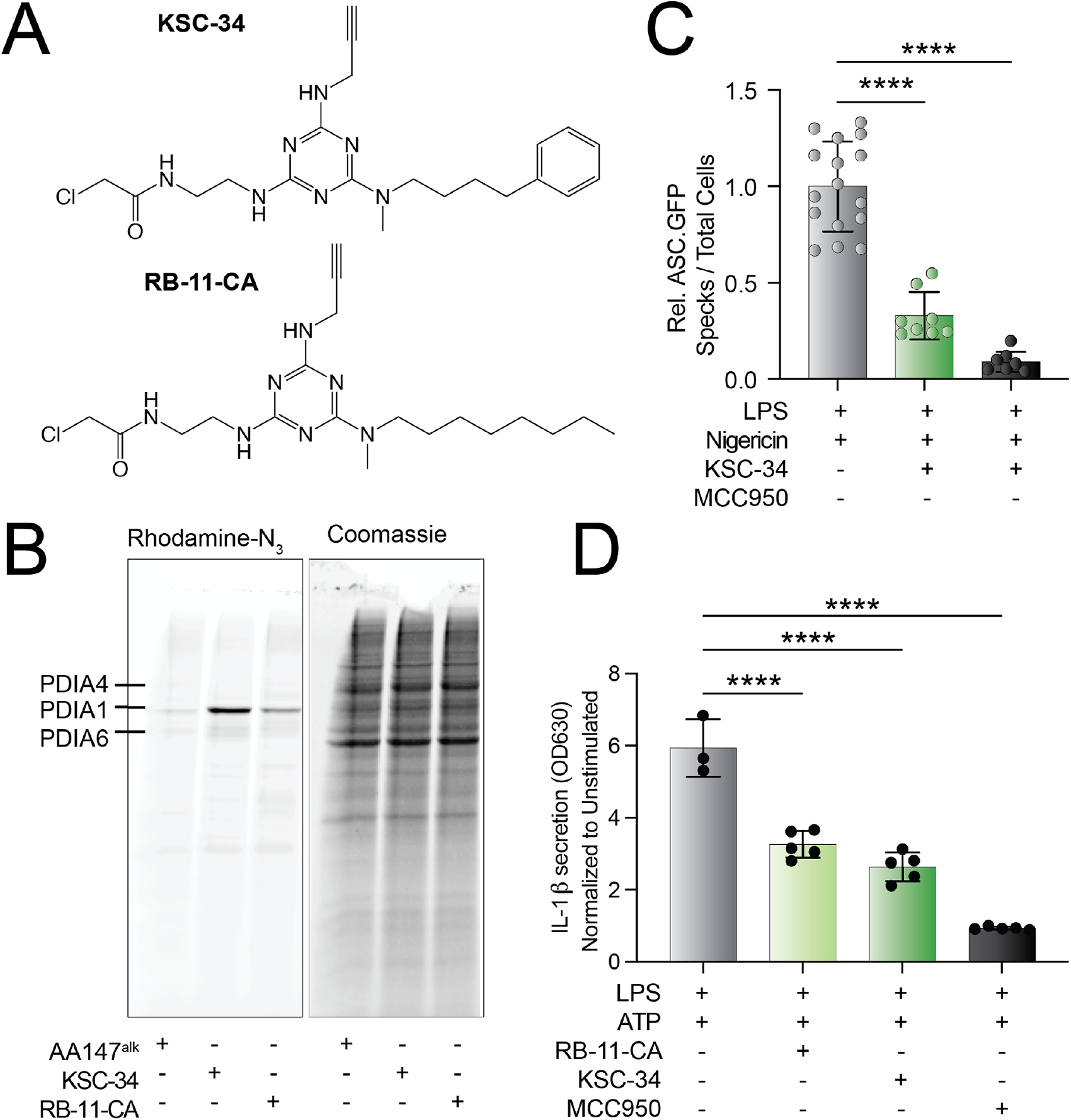
Pharmacologic PDI inhibitors block NLRP3 inflammasome activation. **A.** Chemical structures of the PDI inhibitors KSC-34 and RB-11-CA ^41^. **B**. Relative number of ASC-GFP specks per cell detected using fluorescent microscopy in undifferentiated THP1 ASC-GFP expressing monocyte cells pre-treated with KSC-34 (20μM) and stimulated with LPS (1µg/mL) for 16 h followed by a treatment of nigericin (5 µM) for 3 h. Quantification is relative to the overall average number of specks / total cells across all wells in LPS (1µg/mL) and nigericin (5 µM) stimulated wells. **** p <0.0001 using a Brown-Forsythe and Welch ANOVA test with Dunnett-T3 correction for multiple comparisons. Error bars show SD. **B**. Rhodamine fluorescence and Coomassie staining of an SDS-PAGE gel of lysates from THP1 cells treated for 16 h with AA147^alk^ (10 μM), KSC-34 (20 μM), or RB-11-CA (20 μM) run on an. **C**. Relative number of ASC-GFP specks per cell detected using fluorescent microscopy in undifferentiated THP1 ASC-GFP expressing monocyte cells pre-treated with KSC-34 (20 µM) for 16 h then stimulated with LPS (1µg/mL) for 3 h and nigericin for 3 h (5 μM). **** p <0.0001 using a Brown-Forsythe and Welch ANOVA test with Dunnett-T3 correction for multiple comparisons. Error bars show SD. **D**. Quantification of secreted bioactive IL-1β assessed using the SEAP Quanti-blue assay. HEK Blue IL-1β cells were treated with media conditioned on THP1 monocytes pre-treated with KSC-34 (20 μM) or RB-11-CA (20 μM) and subsequently stimulated for 3 h with LPS (1µg/mL) and 24 h with nigericin (1 μM) relative to an unstimulated, vehicle treated control. * p < 0.05, *** p < 0.001, **** p <0.0001 with one-way ANOVA test with Dunnett correction for multiple comparisons. Error bars show SEM for n=3.

## DISCUSSION

Pharmacologic interventions which suppress the NLRP3 inflammasome activity show significant promise in mitigating the harmful effects of persistent or excessive inflammation.^7,16,17^ Here, we show that the metabolically activated electrophile AA147 can suppress pro-inflammatory IL-1β signaling by inhibiting assembly of the NLRP3 inflammasome. Although AA147 was initially characterized as a preferential activator of the ATF6 arm of the UPR^32^, the inhibition of IL-1β secretion was not dependent on ATF6 signaling. Instead, we show that AA147 impedes inflammasome activity through the covalent modification of PDIs, specifically PDIA1. Genetic ablation of *PDIs*, most notably *PDIA1*, showed similar results to AA147. Importantly, knockdown of *PDIA1* inhibited inflammasome assembly, caspase 1 proteolytic autoactivation, and IL-1β secretion – highlighting the extent to which PDIA1 is involved in NLRP3 inflammasome signaling. Furthermore, selective and structurally distinct inhibitors of PDIA1 similarly reduced assembly of the inflammasome complex as well as IL-1β secretion. Overall, these results indicate that indirectly suppressing inflammasome activation by targeting proteins which regulate NLRP3 inflammasome oligomerization, such as PDIA1, may represent a novel strategy for reducing the pathologic impacts of inflammasome overactivation.

PDIA1 regulates pathologic inflammatory signaling through multiple mechanisms. Previous studies have shown that extracellular PDI is an important regulator of innate immune activity as it can modulate the extent of neutrophil adhesion by impacting the surface expression of molecules including L-selectin^42^ and integrins^43–46^. Blocking antibodies have shown that extracellular PDIs may also be involved in the release of tissue factor and IL-1β from macrophages, although a mechanism for the latter has not been established.^47^ PDIA1 has also been implicated in regulating the secretion of other cytokines, including IL-6 and TNFα.^39^ Here, we used structurally distinct pharmacologic inhibitors of PDIA1 as well as shRNA-mediated silencing to show that PDIA1 can also regulate the processing and release of the potent pro-inflammatory cytokine IL-1β by preventing NLRP3 inflammasome assembly. These studies suggest that PDIA1 activity may be a central regulator of inflammatory signaling across myeloid cell lineages through a diverse range of intracellular and extracellular signaling mechanisms. Therefore, further exploration of the potential of PDI inhibition to mitigate damaging inflammation may present new opportunities to improve patient outcomes across a wide range of disorders.

## METHODS

### Cell culture maintenance and shRNA depletion

The THP1 cells (THP1 null; Invivogen) were a generous gift from the Griffin lab at Scripps Research and the ASC-GFP THP1 (Invivogen) cells were a generous gift from CALIBR. THP1 cell lines were maintained in RPMI 1640 (ATCC formulation, LifeTech cat. A1049101) supplemented with 10% heat-inactivated fetal bovine serum (FBS) (Lifetech cat. 10082147), 100 μg/ml Normocin (Invivogen cat. ant-nr-1) and Pen-Strep (100 μg/ml). Alternating passages of THP1 cells were treated with 200 μg/ml Hygromycin Gold (Invivogen cat. ant-hg-1) as recommended by Invitrogen. ASC-GFP THP1 cells were passaged with 100 μg/ml Zeocin (Fisher Scientific cat. AAJ67140XF). Selection media was fully removed prior to inflammasome stimulation.

THP1 cells were differentiated into macrophages by treating 1M cells /mL with 0.5 μM phorbol 12-myristate 13-acetate (PMA; Sigma; cat P8139) for 3 h on Day 0. After the initial media change following PMA, media was refreshed daily for two additional days. On Day 2, compounds were added to media for pre-treatment. The following day, THP1 derived macrophages were stimulated with inflammasome priming and activation stimuli; compound was readded to media with each media change after initial pre-treatment.

For shRNA depletion, viruses expressing specific shRNAs were prepared as previously described.38 Briefly, one 10 cm dish of HEK293T cells per shRNA was transiently transfected with 8 μg shRNA construct, 4 μg REV (pRSV-rev), 4 μg RRE (pMDL-RRE), and 4 μg VSV-G (pMD2.G). Transfection reagents were removed after a 24 h incubation, followed by a 24 h incubation for viral production in THP1 cell media. A 1:1 ratio of virus-containing media and fresh media was added to THP1 cells or ASC-GFP THP1 cells for 24 h. Transfected cells were puromycin-selected (5 μg/L) (Sigma-Aldrich; cat P8833) for 7 days. Knockdown was confirmed by real-time quantitative polymerase chain reaction (RT-qPCR) and immunoblotting.

IL-1β Reporter HEK 293 cells were a kind gift from CALIBR and were cultured in DMEM supplemented with 10% heat inactivated FBS (Lifetech cat. 10082147), 2 mM glutamine and Pen-Strep (100 μg/ml). Cells were subcultured in 100 μg/ml of Zeocin as recommended by manufacturer. Trypsin was not used to detach cells, as it can interfere with the secreted alkaline phosphatase (SEAP) reporter. HEK293T cells were cultured in high glucose DMEM (4.5 g/L) (Corning cat. 15-017-CV) supplemented with 10% FBS and Pen-Strep (100 μg/ml).

### Compounds and Treatments

Ultra-pure TLR4 specific lipopolysaccharide derived from *E. coli* strain 0111:B4 (LPS; Invivogen cat. tlrl-3pelps) at a dose of 1µg/mL was used to induce NF-κB transcription activity for inflammasome priming across all experiments. Inflammasome assembly was induced by administering cells with 5mM ATP (Jena Biosciences cat. NU-1010) or nigericin (Invivogen cat. tlrl-nig) with dose, as indicated.

AA147 and AA147^alk^ were previously reported^33^ and kind gifts from the Kelly Lab at Scripps Research, suspended in dimethyl sulfoxide (DMSO), and administered at 10 μM. PF429242 (Sigma-Aldrich; cat. SML0667) was resuspended in water and administered at 10 μM. CP7 was obtained from the Walter Lab at UCSF, resuspended in DMSO, and administered at 6 μM. RB-11-CA and KSC-34 were previously reported^41^ and were kind gifts from the Weerapana Lab at Boston College. Compounds were resuspended in DMSO and administered at 20 μM. MCC950 (Selleck Chem; cat. S7809) was resuspended in water and administered at 1 μM concurrently with the addition of LPS for all experiments. Z-VAD-FMK was administered at 50 μM at the same time as the secondary stimulus.

### Plasmids and Antibodies

For viral transfection, the following plasmids were used: REV (pRSV-rev; Addgene cat. 12253), RRE (pMDL-RRE; Addgene cat. 12251), and VSV-G (pMD2. G; Addgene cat. 12259). Mission shRNA plasmids in pLKO.1 vectors were obtained from La Jolla Institute for Allergy and Immunology (LJI) and were previously used in Paxman et al.38 The specific target sequences for viral plasmid of these shRNAs are below:

Target (Designation) shRNA sequence

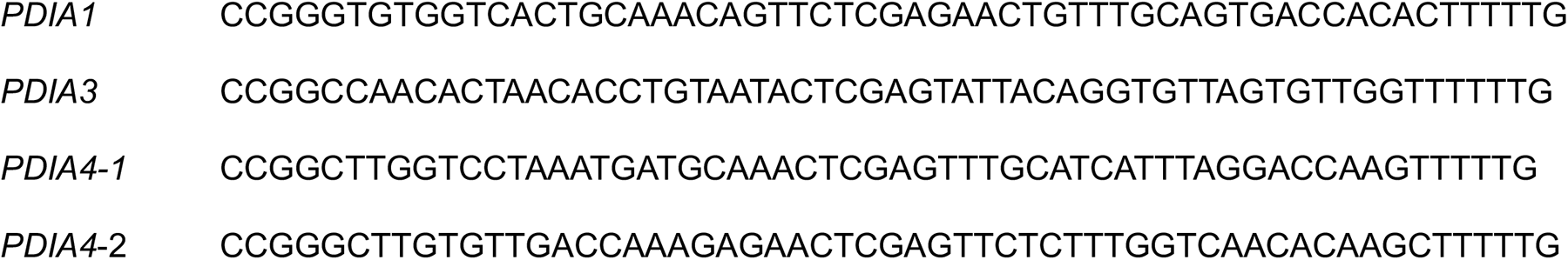
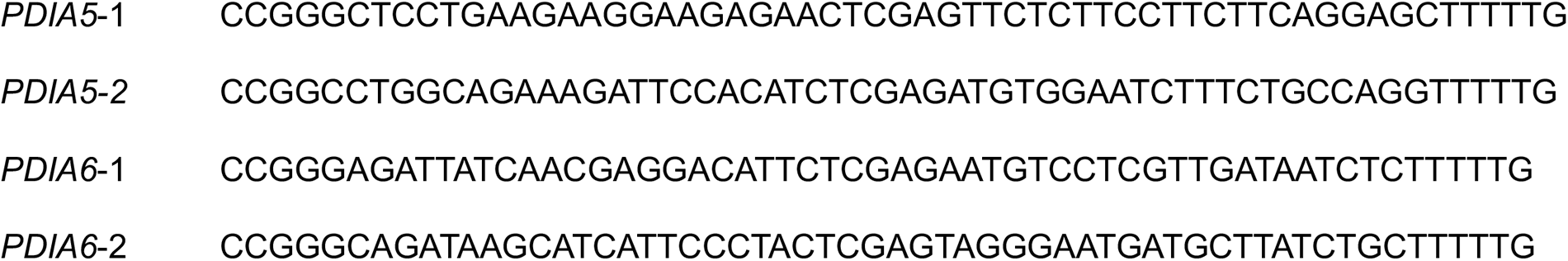

The following primary antibodies were diluted as noted in 5% bovine serum albumin (BSA) in TBS and incubated overnight: KDEL (1:1000; Enzo cat. ADI-SPA-827-F), NLRP3 (1:1000; Cell Signaling cat. 15101), CASP1 (1:1000; Cell Signaling cat. 3866), ASC (1:1000, Cell Signaling cat. 13833), IL-1β (1:1000, GeneTex cat. GTX74034), PDIA1 (1:1000, Cell Signaling cat. 45596), Tubulin (1:2000; Sigma-Aldrich T6074), GAPDH (1:1000, GeneTex cat. GTX627408), NEK7 (1:1000, Cell Signaling cat. 3057).

### Inflammasome assembly assay

ASC. GFP THP1 cells were plated at a density of 20,000 cells / well in a 384 well plate and treated with compounds for 16 h. Cells were then primed with LPS (1µg/mL) for either 3 h or 16 h as noted. Subsequently, cells were stimulated with nigericin (10 μM) for 3 hours. Nuclei were stained using 5 µg/mL Hoechst 33342 (Thermo cat. H3570). Images were captured using Cellomics Cell Insight imaging reader (Thermo). An air ×10 lens was used to capture one image per well. For image analysis, stained nuclei were enumerated as representing the total number of live cells captured. ASC-GFP speck formation was calculated using an algorithm within the Cell Insight software, which factors both signal intensity and area size. Numerical results from the analyzed images were later exported for analysis using Prism 9 (GraphPad, San Diego, CA)

### IL-1β assays

Undifferentiated THP1 cells were plated in fresh media at a density of 750K/mL and compounds were concurrently administered, as noted. The following day, cells were stimulated using LPS (1µg/mL) for 3 h and then ATP (5 mM) or nigericin (1 μM) were added directly to the cells. Cells were incubated at 37o for 24 h following the addition of the secondary stimulus. Cells and conditioned media were centrifuged at a low speed to remove suspended monocytes. The supernatant was added IL-1β Reporter HEK 293 Cells (Invivogen cat. hkb-il1bv2) following the manufacturer’s protocol. Quanti-blue (Invivogen cat. rep-qbs) was used to detect the levels of the secreted alkaline phosphatase reporter following manufacturer’s protocol. Absorbance was measured using SPECTRAmax PLUS 384 (Molecular Devices) plate reader at OD630.

For assays performed on THP1 derived macrophages, macrophages were treated on day 3 for 3 h with Ultrapure TLR4 specific lipopolysaccharide derived from E. coli strain 0111:B4 (LPS; Invivogen cat. tlrl-3pelps). This media was changed and ATP (5 mM) (Jena Biosciences cat. NU-1010) was added for 24 h prior to collection of conditioned media. Conditioned media was added directly to the IL-1β Reporter HEK 293 cells. Media for the ELISA was immediately stored after collection at -80°C for subsequent testing in the enzyme-linked immunoassay (R&D systems). The Human IL-1β Quantikine ELISA Kit (R&D Systems cat. DLB50) was used to quantify levels of IL-1β according to the manufacturer’s protocol. RAW264.7 cells were plated in the morning and pre-treated in the afternoon on the same day as plating. The following day cells stimulated for 3 h with Ultrapure TLR4 specific lipopolysaccharide derived from E. coli strain 0111:B4 (LPS; Invivogen cat. tlrl-3pelps). This media was changed and ATP (5 mM) (Jena Biosciences cat. NU-1010) was added for 24 h prior to collection of conditioned media. IL-1β levels were detected in conditioned media using SDS-PAGE followed by immunoblotting.

### ASC-GFP Flow Cytometry

ASC-GFP THP1 cells were plated at a density of 750K cells/mL were plated in 100 DL in a 96 well plate. Cells were pre-treated for 16 h with AA147. The following day, cells were stimulated with LPS (1µg/mL) for 3 h and immediately analyzed via flow cytometry on a NovoCyte 3000 (Acea). GFP fluorescence was detected using ex. 488 nm, em. 530/30 nm channel. Analysis and gating were performed using FlowJo software (BD Biosciences, San Diego). A minimum of 10,000 cells per sample were used to calculate the geometric mean GFP signal.

### Quantitative reverse-transcriptase polymerase chain reaction (qRT-PCR)

Cells were rinsed with PBS, lysed, and total RNA was collected using the QuickRNA mini kit (Zymo) according to the manufacturer’s instructions. The relative quantification of mRNA per sample was calculated using qPCR with reverse transcription (RT-qPCR) of the endogenous control RPLP2 (RiboP). RNA yield was quantified using Nanodrop. cDNA was generated from 300 ng of RNA using High-Capacity cDNA Reverse Transcription Kit (Advanced Biosystems; cat. 4368814). qPCR reactions were prepared using Power SYBR Green PCR Master Mix (Applied Biosystems; cat. 4367659), and primers were obtained from Integrated DNA Technologies. Amplification reactions were run in an ABI 7900HT Fast Real Time PCR machine with an initial melting period of 95 °C for 5 min and then 45 cycles of 10 s at 95 °C, 30 s at 60 °C.

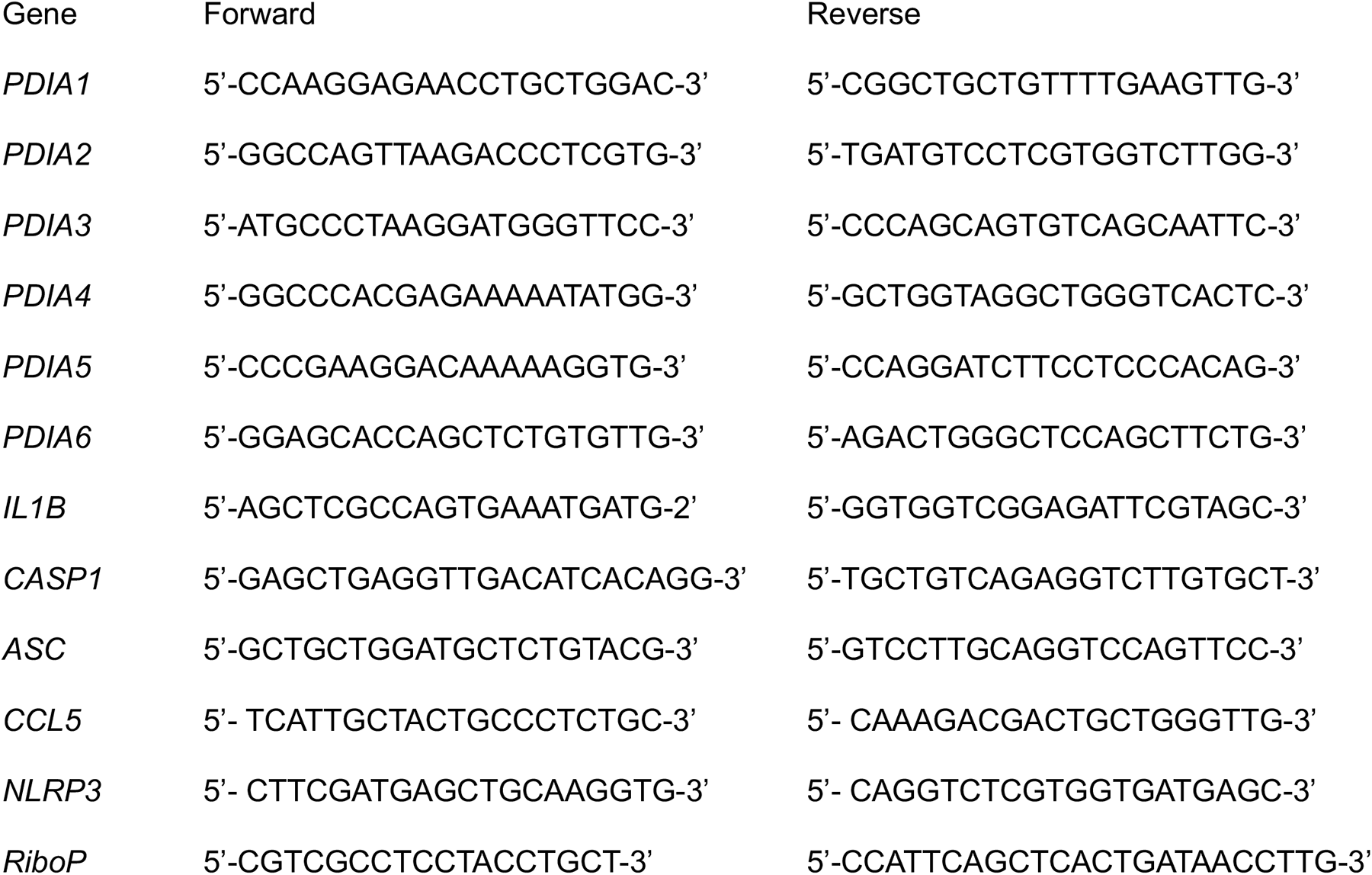

### Bioluminescent caspase 1 cleavage assay

The caspase 1 glo (Promega; cat. G9951) was followed according to manufacturer’s instructions. Briefly, undifferentiated THP1 cells were plated at a density of 750K cells/mL and treated for 16 h with LPS (1μg/mL) in a flask. The following day, cells were removed from LPS media, resuspended in fresh media at a density of 1M cells /mL, and 25 μL of the suspension was plated in a 384 well dish. 20 μM Nigericin was added to individual wells for 45 minutes. Subsequently, caspase 1 reagent as well as the proteasome inhibitor MG132, and ac-YVAD were added to respective treatments and or controls. Reactions were incubated in the dark at room temperature for a minimum of 1 prior to luminescence being read on a Tecan F200 Pro microplate reader.

### Rhodamine labeling

Rhodamine-azide labeling reactions were performed in a 50 μL reaction containing 1.5 μg of protein from lysate from THP1 cell lysate treated with respective compounds for 16 h. Click reactions contained a final concentration of 0.8 mM CuSO4, 1.6 mM BTTAA (Sigma cat. 906328), 5mM Sodium Ascorbate (Sigma cat. A4034), and 0.1 mM TAMRA azide (Santa Cruz cat. sc-482005). Reactions were incubated at 4oC overnight, then denatured with 1 × Laemmli buffer + 100 mM dithiothreitol (DTT) and boiled before being separated by SDS-PAGE. Gel was then stained with 0.1% Coomassie Blue R250 in 10% acetic acid/50% methanol solution. Both rhodamine fluorescence and Coomassie staining were imaged on a ChemiDoc Imaging Systems (BioRad).

### NLRP3 labeling and immunopurification

THP-1 cells were treated with either KSC-34 or RB-11-CA for 16 h. Cell lysates were prepared via sonication in PBS. Biotin labeling reactions were performed using 1.7 mM TBTA, 50 mM CuSO4, 5 mM Biotin-PEG3-azide, and 50 mM tris(2-carboxyethyl)phosphine (TCEP). The protein was purified using MeOH precipitation, resuspended in 0.1% SDS in PBS and streptavidin agarose slurry added. Streptavidin enrichment and elution was performed as previously described.57

### SDS-Page and Immunoblotting

Briefly, cells were lysed in lysis buffer (50 mM Tris, pH 7.5, 150 mM NaCl, 20 mM Hepes pH 7.4, 1 mM EDTA, 1% Triton X-100, and protease inhibitor cocktail (Pierce cat. A32955)). The total protein concentration in cellular lysates was normalized using the Bio-Rad protein assay. Lysates were then denatured with 1 × Laemmli buffer + 100 mM dithiothreitol (DTT) and boiled before being separated by SDS-PAGE. Samples were transferred onto nitrocellulose membranes (Bio-Rad cat. 1620112). Membranes were then incubated overnight at 4 °C with primary antibodies with dilutions, as noted. Membranes were washed in TBS-T, incubated with the species-appropriate IR-Dye conjugated secondary antibodies diluted at 1:10,000, and analyzed using the Odyssey Infrared Imaging System (LI-COR Biosciences). Quantification was carried out with LI-COR Image Studio software.

## ACKNOWLEDGEMENTS

We thank John Griffin and Laura Healy at Scripps Research for experimental advice related to the work described in this manuscript. This work was funded by the National Institutes of Health (DK107604 to RLW).

## AUTHOR CONTRIBUTION

JR and CR conceived, designed, performed, and interpreted the experiments. MB and RLW conceived and designed experiments and supervised the project. JR wrote the manuscript. CR, MB, and RLW revised the manuscript and provided approval for submission.

---

**Figure S1.**
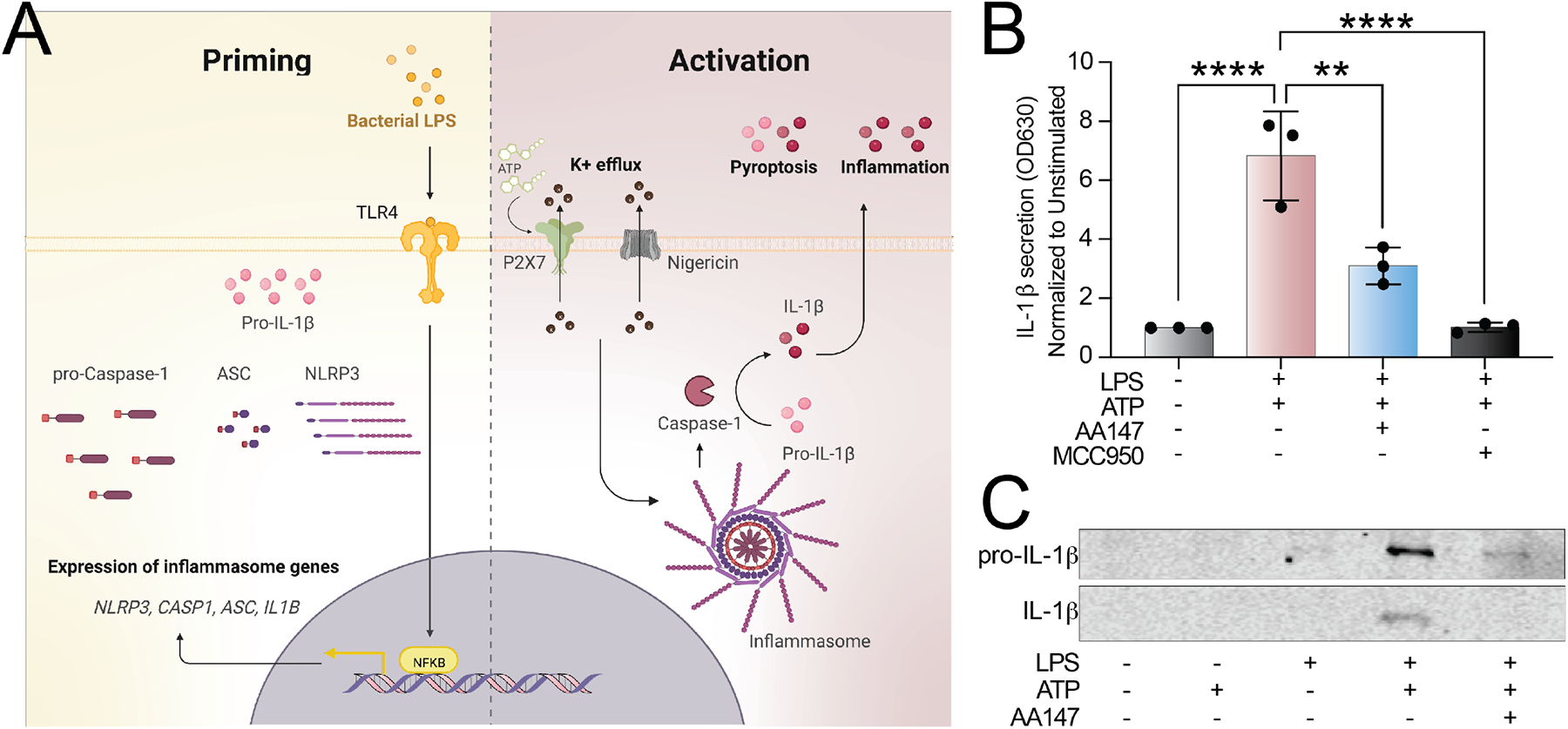
AA147 reduces NLRP3 inflammasome assembly and IL-1β secretion. **A**. Illustration of NLRP3 priming and activation steps required for activity prepared in BioRender. **B.** Quantification of secreted bioactive IL-1β assessed using the SEAP Quanti-blue assay. HEK Blue IL-1β cells were treated with media conditioned on THP1 monocyte-derived macrophages pre-treated with AA147 (10 µM) or vehicle for 16 h then stimulated with LPS (1µg/mL) for 3 h and incubated with ATP (5 mM) for 24 h relative to an unstimulated, vehicle treated control. **** p <0.0001, *** p < 0.001, ** p<0.01 for one-way ANOVA test with Dunnett correction for multiple comparisons. Error bars show SEM for n=3. **C.** Immunoblot of conditioned media from RAW264.7 mouse macrophage cells pre-treated with AA147 (10 µM) or vehicle for 16 h and then stimulated with LPS (1µg/mL) for 3 h and ATP (5 mM) for 24 h.

**Figure S2.**
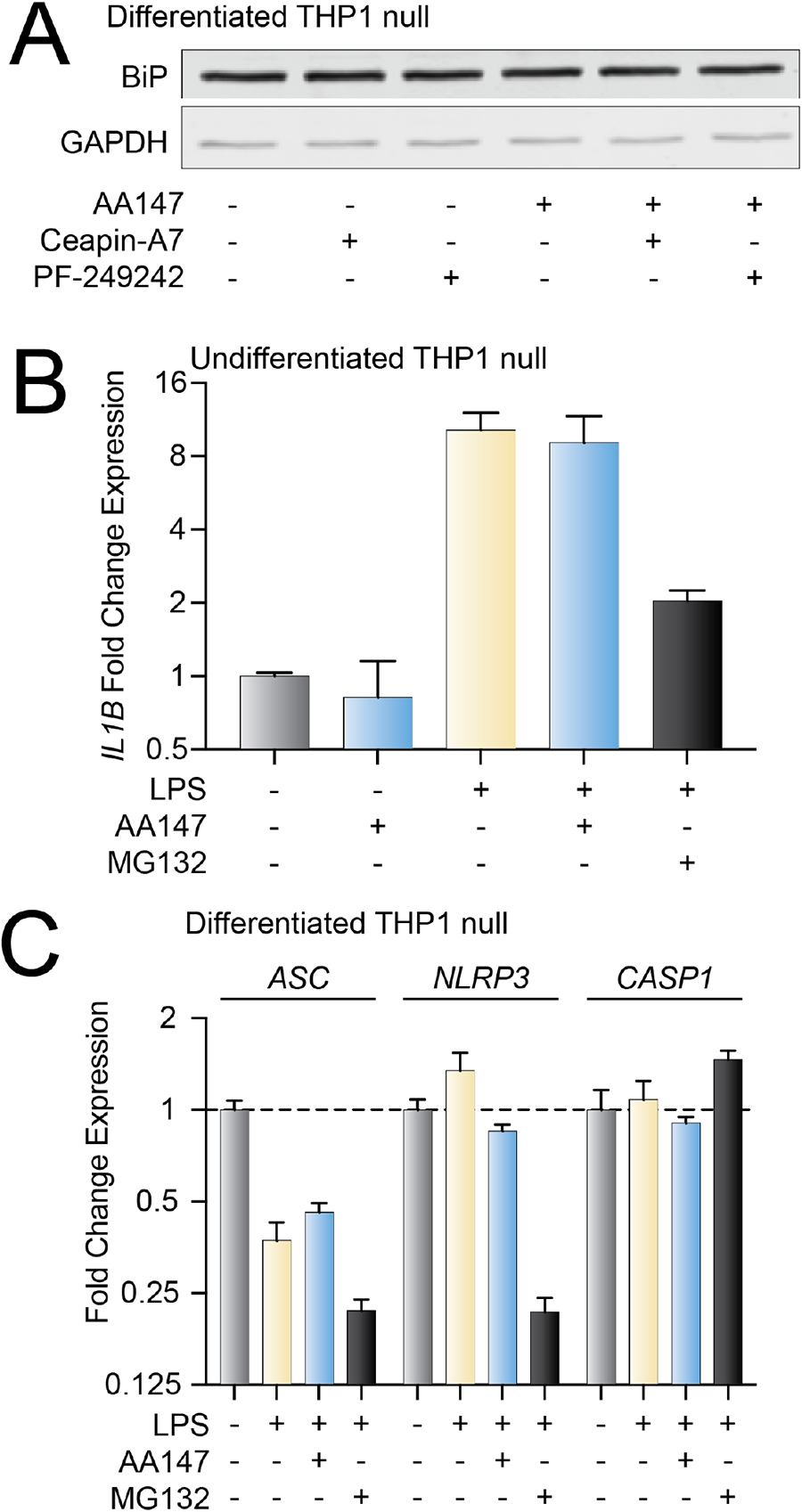
AA147-dependent inhibition of NLRP3 inflammasomes is independent of ATF6 activation. **A**. Immunoblot from THP1 monocyte-derived macrophages treated for 16 h with vehicle or AA147 (10 µM) in the presence or absence of the ATF6 inhibitor Ceapin-7 (6 µM). **B.** mRNA expression measured using qPCR, of the pro-inflammatory cytokine *IL1B* in THP1 monocyte-derived macrophages treated for 16 h with vehicle or AA147 (10 µM) and subsequently stimulated with LPS (1µg/mL) for 3 h. Error bars show 95% CI. **C.** mRNA expression measured using qPCR, of the inflammasome component *ASC*, *CASP1*, and *NLRP3* in THP1 monocyte-derived macrophages treated for 16 h with vehicle or AA147 (10 µM) and subsequently stimulated with LPS (1µg/mL) for 3 h. Error bars show 95% CI.

**Figure S3.**
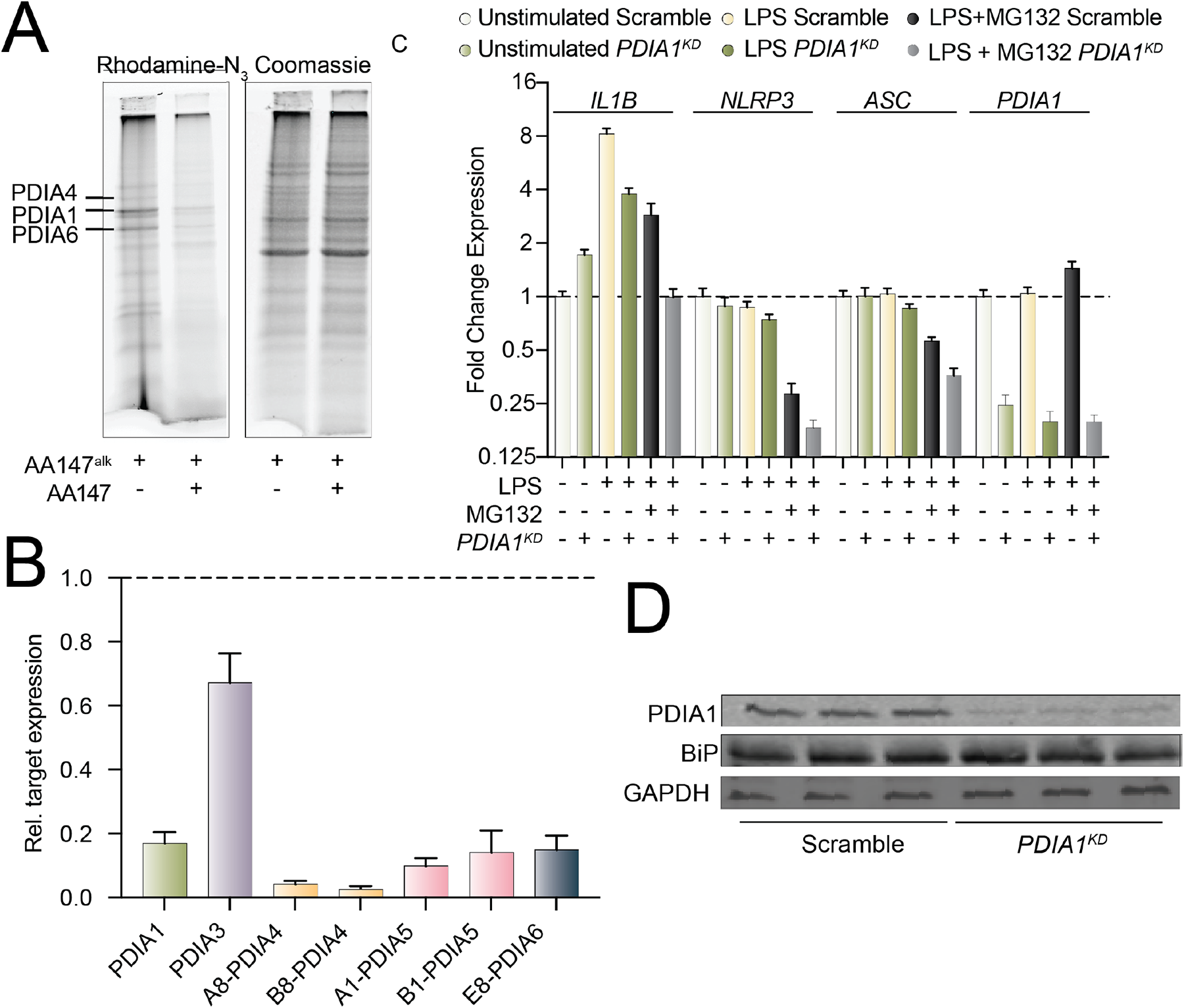
*PDIA1* depletion inhibits NLRP3 inflammasome activation. **A**. THP1 cells were treated with AA147^alk^ (10 μM) for 16 h, alone and in competition with AA147 (50 μM). Rhodamine fluorescence and Coomassie stain of SDS-PAGE gel of lysates. **B.** qPCR of THP1 cells stably expressing an shRNA targeted toward the respective PDI. Error bars show 95% CI. **C**. mRNA expression of the inflammasome component *IL1B*, *ASC*, and *NLRP3* and *PDIA1* measured by qPCR in THP1 monocyte-derived macrophages stimulated with LPS (1µg/mL) for 3 h. Error bars show 95% CI. **D.** Immunoblot of cell lysates from PDIA1^KD^ or scramble THP1 cells.

**Figure S4.**
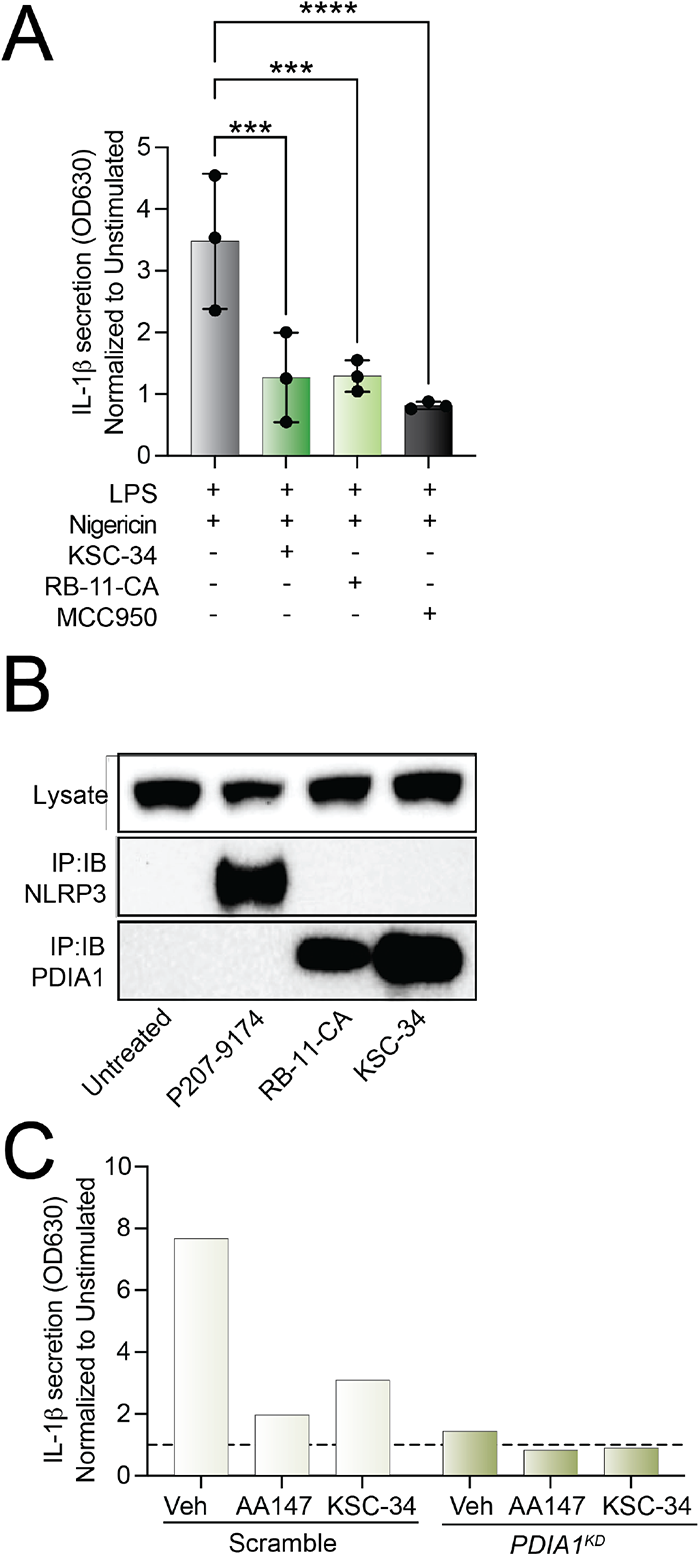
Pharmacologic PDI inhibitors block NLRP3 inflammasome activation. **A.** Quantification of secreted bioactive IL-1β assessed using the SEAP Quanti-blue assay. HEK Blue IL-1β cells treated with media conditioned on THP1 monocytes pre-treated with AA147 (10 µM), KSC-34 (20 μM), or RB-11-CA (20 μM) and subsequently stimulated for 3 h with LPS (1µg/mL) and 24 h with nigericin (1 μM) relative to an unstimulated, vehicle treated control. * p < 0.05, *** p < 0.001, **** p <0.0001 with one-way ANOVA test with Dunnett correction for multiple comparisons. Error bars show SEM for n=3. **B.** Biotin-Streptavidin immunoprecipitation and immunoblotting of cellular lysates from THP1 cells treated with RB-11-CA (20 µM) or KSC-34 (20 µM) for 16 h then subsequently conjugated to a biotin tag. D053-0367 is an unpublished positive control. **C**. Quantification of secreted bioactive IL-1β assessed using the SEAP Quanti-blue assay. HEK Blue IL-1β cells treated with media conditioned on THP1 monocytes stably expressing either a PDIA1 shRNA or a scramble shRNA. THP1 cells were pre-treated with AA147 (10 μM) or KSC-34 (20 μM) for 16 h, then stimulated for 3 h with LPS (1 µg/mL) and 24 h with nigericin (1 μM). Quantification is relative to an unstimulated scramble-shRNA expressing vehicle treated control.

## REFERENCES

1 Lamkanfi, M. & Dixit, V. M. Mechanisms and functions of inflammasomes. Cell 157, 1013–1022 (2014). 10.1016/j.cell.2014.04.007

2 Gross, O., Thomas, C. J., Guarda, G. & Tschopp, J. The inflammasome: an integrated view. Immunological Reviews 243, 136–151 (2011). 10.1111/j.1600-065X.2011.01046.x

3 Kelley, N., Jeltema, D., Duan, Y. & He, Y. The NLRP3 Inflammasome: An Overview of Mechanisms of Activation and Regulation. Int J Mol Sci 20, 3328 (2019). 10.3390/ijms20133328

4 Broz, P. & Dixit, V. M. Inflammasomes: mechanism of assembly, regulation and signalling. Nature Reviews Immunology 16, 407–420 (2016). 10.1038/nri.2016.58

5 Netea, M. G. et al. Differential requirement for the activation of the inflammasome for processing and release of IL-1β in monocytes and macrophages. Blood 113, 2324–2335 (2009). 10.1182/blood-2008-03-146720

6 Sharif, H. et al. Structural mechanism for NEK7-licensed activation of NLRP3 inflammasome. Nature 570, 338–343 (2019). 10.1038/s41586-019-1295-z

7 Swanson, K. V., Deng, M. & Ting, J. P. Y. The NLRP3 inflammasome: molecular activation and regulation to therapeutics. Nature Reviews Immunology 19, 477–489 (2019). 10.1038/s41577-019-0165-0

8 Tong, Y. et al. NLRP3 Inflammasome and Its Central Role in the Cardiovascular Diseases. Oxidative medicine and cellular longevity 2020, 4293206 (2020). 10.1155/2020/4293206

9 Dinarello, C. A. Interleukin-1 in the pathogenesis and treatment of inflammatory diseases. Blood 117, 3720–3732 (2011). 10.1182/blood-2010-07-273417

10 Aday, A. W. & Ridker, P. M. Antiinflammatory Therapy in Clinical Care: The CANTOS Trial and Beyond. Frontiers in cardiovascular medicine 5, 62 (2018). 10.3389/fcvm.2018.00062

11 Aksentijevich, I. et al. The clinical continuum of cryopyrinopathies: Novel CIAS1 mutations in North American patients and a new cryopyrin model. Arthritis & Rheumatism 56, 1273–1285 (2007). 10.1002/art.22491

12 Brydges, S. D. et al. Inflammasome-Mediated Disease Animal Models Reveal Roles for Innate but Not Adaptive Immunity. Immunity 30, 875–887 (2009). 10.1016/j.immuni.2009.05.005

13 Mikkelsen, R. R. et al. Immunomodulatory and immunosuppressive therapies in cardiovascular disease and type 2 diabetes mellitus: A bedside-to-bench approach. European Journal of Pharmacology 925, 174998 (2022). 10.1016/j.ejphar.2022.174998

14 Larsen, C. M. et al. Interleukin-1–Receptor Antagonist in Type 2 Diabetes Mellitus. New England Journal of Medicine 356, 1517–1526 (2007). 10.1056/NEJMoa065213

15 Ismael, S., Zhao, L., Nasoohi, S. & Ishrat, T. Inhibition of the NLRP3-inflammasome as a potential approach for neuroprotection after stroke. Scientific Reports 8, 5971 (2018). 10.1038/s41598-018-24350-x

16 Abbate, A. et al. Interleukin-1 and the Inflammasome as Therapeutic Targets in Cardiovascular Disease. Circulation Research 126, 1260–1280 (2020). doi:10.1161/CIRCRESAHA.120.315937

17 Mangan, M. S. J. et al. Targeting the NLRP3 inflammasome in inflammatory diseases. Nat Rev Drug Discov 17, 588–606 (2018). 10.1038/nrd.2018.97

18 Yang, Y., Wang, H., Kouadir, M., Song, H. & Shi, F. Recent advances in the mechanisms of NLRP3 inflammasome activation and its inhibitors. Cell Death Dis 10, 128 (2019). 10.1038/s41419-019-1413-8

19 Li, Y. et al. Inflammasomes as therapeutic targets in human diseases. Signal Transduction and Targeted Therapy 6, 247 (2021). 10.1038/s41392-021-00650-z

20 Lythgoe, M. P. & Prasad, V. Repositioning canakinumab for non-small cell lung cancer—important lessons for drug repurposing in oncology. British Journal of Cancer (2022). 10.1038/s41416-022-01893-5

21 Ridker, P. M. et al. Antiinflammatory Therapy with Canakinumab for Atherosclerotic Disease. New England Journal of Medicine 377, 1119–1131 (2017). 10.1056/NEJMoa1707914

22 Coll, R. C. et al. MCC950 directly targets the NLRP3 ATP-hydrolysis motif for inflammasome inhibition. Nat Chem Biol 15, 556–559 (2019). 10.1038/s41589-019-0277-7

23 Coll, R. C. et al. A small-molecule inhibitor of the NLRP3 inflammasome for the treatment of inflammatory diseases. Nat Med 21, 248–255 (2015). 10.1038/nm.3806

24 van Hout, G. P. et al. The selective NLRP3-inflammasome inhibitor MCC950 reduces infarct size and preserves cardiac function in a pig model of myocardial infarction. Eur Heart J 38, 828–836 (2017). 10.1093/eurheartj/ehw247

25 Primiano, M. J. et al. Efficacy and Pharmacology of the NLRP3 Inflammasome Inhibitor CP-456,773 (CRID3) in Murine Models of Dermal and Pulmonary Inflammation. The Journal of Immunology 197, 2421–2433 (2016). 10.4049/jimmunol.1600035

26 Marchetti, C. et al. OLT1177, a β-sulfonyl nitrile compound, safe in humans, inhibits the NLRP3 inflammasome and reverses the metabolic cost of inflammation. Proceedings of the National Academy of Sciences 115, E1530–E1539 (2018). doi:10.1073/pnas.1716095115

27 Klück, V. et al. Dapansutrile, an oral selective NLRP3 inflammasome inhibitor, for treatment of gout flares: an open-label, dose-adaptive, proof-of-concept, phase 2a trial. Lancet Rheumatol 2, e270–e280 (2020). 10.1016/s2665-9913(20)30065-5

28 He, H. et al. Oridonin is a covalent NLRP3 inhibitor with strong anti-inflammasome activity. Nature Communications 9, 2550 (2018). 10.1038/s41467-018-04947-6

29 Chen, Y. et al. RRx-001 ameliorates inflammatory diseases by acting as a potent covalent NLRP3 inhibitor. Cell Mol Immunol 18, 1425–1436 (2021). 10.1038/s41423-021-00683-y

30 Hooftman, A. et al. The Immunomodulatory Metabolite Itaconate Modifies NLRP3 and Inhibits Inflammasome Activation. Cell Metab 32, 468–478.e467 (2020). 10.1016/j.cmet.2020.07.016

31. Stanton, C., et al. Covalent targeting as a common mechanism for inhibiting NLRP3 inflammasome assembly. bioRxiv (2023). 10.1101/2023.06.01.543248

32 Plate, L. et al. Small molecule proteostasis regulators that reprogram the ER to reduce extracellular protein aggregation. eLife 5 (2016). 10.7554/eLife.15550

33 Paxman, R. et al. Pharmacologic ATF6 activating compounds are metabolically activated to selectively modify endoplasmic reticulum proteins. eLife 7 (2018). 10.7554/eLife.37168

34 Rius, B. et al. Pharmacologic targeting of plasma cell endoplasmic reticulum proteostasis to reduce amyloidogenic light chain secretion. Blood Advances 5, 1037–1049 (2021). 10.1182/bloodadvances.2020002813

35 Zhou, R., Yazdi, A. S., Menu, P. & Tschopp, J. A role for mitochondria in NLRP3 inflammasome activation. Nature 469, 221–225 (2011). 10.1038/nature09663

36 Rosarda, J. D. et al. Metabolically Activated Proteostasis Regulators Protect against Glutamate Toxicity by Activating NRF2. ACS Chemical Biology 16, 2852–2863 (2021). 10.1021/acschembio.1c00810

37 Gallagher, C. M. & Walter, P. Ceapins inhibit ATF6alpha signaling by selectively preventing transport of ATF6alpha to the Golgi apparatus during ER stress. eLife 5 (2016). 10.7554/eLife.11880

38 Hawkins, J. L. et al. Pharmacologic inhibition of site 1 protease activity inhibits sterol regulatory element-binding protein processing and reduces lipogenic enzyme gene expression and lipid synthesis in cultured cells and experimental animals. The Journal of pharmacology and experimental therapeutics 326, 801–808 (2008). 10.1124/jpet.108.139626

39 Xiao, Y. et al. Protein Disulfide Isomerase Silence Inhibits Inflammatory Functions of Macrophages by Suppressing Reactive Oxygen Species and NF-κB Pathway. Inflammation 41, 614–625 (2018). 10.1007/s10753-017-0717-z

40 O’Brien, M. et al. A bioluminescent caspase-1 activity assay rapidly monitors inflammasome activation in cells. Journal of Immunological Methods 447, 1–13 (2017). 10.1016/j.jim.2017.03.004

41 Cole, K. S. et al. Characterization of an A-Site Selective Protein Disulfide Isomerase A1 Inhibitor. Biochemistry 57, 2035–2043 (2018). 10.1021/acs.biochem.8b00178

42 Bennett, T. A., Edwards, B. S., Sklar, L. A. & Rogelj, S. Sulfhydryl Regulation of L-Selectin Shedding: Phenylarsine Oxide Promotes Activation-Independent L-Selectin Shedding from Leukocytes1. The Journal of Immunology 164, 4120–4129 (2000). 10.4049/jimmunol.164.8.4120

43 Cho, J. et al. Protein disulfide isomerase capture during thrombus formation in vivo depends on the presence of β3 integrins. Blood 120, 647–655 (2012). 10.1182/blood-2011-08-372532

44 Hahm, E. et al. Extracellular protein disulfide isomerase regulates ligand-binding activity of αMβ2 integrin and neutrophil recruitment during vascular inflammation. Blood 121, 3789–3800 (2013). 10.1182/blood-2012-11-467985

45 Kim, K. et al. Platelet protein disulfide isomerase is required for thrombus formation but not for hemostasis in mice. Blood 122, 1052–1061 (2013). 10.1182/blood-2013-03-492504

46 Lahav, J. et al. Sustained integrin ligation involves extracellular free sulfhydryls and enzymatically catalyzed disulfide exchange. Blood 100, 2472–2478 (2002). 10.1182/blood-2001-12-0339

47 Furlan-Freguia, C., Marchese, P., Gruber, A., Ruggeri, Z. M. & Ruf, W. P2X7 receptor signaling contributes to tissue factor-dependent thrombosis in mice. J Clin Invest 121, 2932–2944 (2011). 10.1172/jci46129

